# *In Vivo* Rescue of Cardiomyopathy Through Cardiac Prime Editing

**DOI:** 10.64898/2026.02.10.705148

**Authors:** Wenjing Liang, Lindsey M. Rollosson, Emilee Easter, Huanyu Zhou, Cristina Dee-Hoskins, Amara Greer-Short, Timothy Hoey, Laura M. Lombardi, Kathryn N. Ivey, Ze Cheng

## Abstract

Heart disease affects millions of individuals and prime editing (PE) may enable curative therapies that address the underlying drivers of heart disease. Here we describe the establishment and optimization of an *in vivo* cardiac PE platform which mediates efficient editing in the heart with no detectable editing in the liver. We performed a proof-of-concept test on RNA binding motif protein 20 *(RBM20)*, which if mutated, can cause dilated cardiomyopathy (DCM) in humans. Our dual-AAV based PE therapeutic rescued cardiomyopathy phenotypes in the heterozygous *Rbm20^R636Q^* mouse model. To further develop PE targeting human *RBM20*, we introduced a novel humanized mouse model carrying human *RBM20* wildtype (WT) or R634Q mutant sequences and displaying *RBM20* cardiomyopathy phenotypes. Our human *RBM20* PE therapeutic efficiently corrected the pathogenic mutation and rescued phenotypes in the humanized *RBM20* mouse model. Our findings demonstrate the potential of *in vivo* cardiac PE in treating heart disease, offer a valuable humanized DCM mouse model for developing various therapies, and present an optimized *in vivo* PE platform that can be adopted for targeting other organs and tissues.

## INTRODUCTION

Heart disease is the leading cause of death in the world, and the unmet need for curative therapies that target the underlying causes of heart disease is high(*1*). Heart disease comes in many forms, affects individuals at many ages, and can be a result of many factors, including cardiomyocyte malfunction caused by genetic mutations(*2*).

Gene editing technologies may correct the underlying genetic causes of cardiac conditions and base editing (BE) has been developed for treating heart disease in mouse models(*3–9*), but its therapeutic application is restricted by limited conversion types, the necessity for a nearby PAM sequence, and unintended bystander editing(*3, 10*). In contrast, PE has the potential to be a transformative therapeutic platform for treating genetic cardiomyopathies due to its ability to mediate precise targeted insertions, deletions, and all 12 possible classes of point mutations in a relatively wide editing window(*10–12*), however, *in vivo* PE technology has not previously been engineered for efficient and selective editing in cardiomyocytes following systemic administration and the efficacy of *in vivo* PE in treating heart disease has not previously been demonstrated.

PE utilizes a fusion protein machinery consisting of a programmable nickase and an engineered reverse transcriptase, guided by a prime editing guide RNA (pegRNA) to the target for installation of the desired edit(*11*). Optionally, a second guide RNA, the nicking guide RNA (ngRNA), can be included in the PE system to enhance editing efficiency(*11*). For PE to be translated into cardiac therapeutics that correct pathogenic mutations *in vivo* in cardiomyocytes following systemic administration, methods of arranging PE components in a clinically validated delivery system and efficiently targeting cardiomyocytes are needed. Previous reports introduced dual-AAV PE designs and demonstrated editing and/or efficacy of *in vivo* PE in the liver, the brain, or the retina in relevant mouse models(*13–20*). Among these studies, Davis et al. and Wei et al. performed systemic administration and showed that v3em PE-AAV system and split-PE-367 system efficiently edited the liver, while the editing efficiency in the heart was suboptimal(*15, 17*).

*RBM20* encodes a splicing regulator and mutations in *RBM20* can cause DCM, accounting for 2% to 6% of cases of familial DCM and displaying high penetrance and aggressive progression with high rates of heart failure, arrhythmias, and sudden cardiac death(*21, 22*). Many *RBM20* disease-causing mutations cluster in a small arginine/serine-rich region(*21, 22*), which may enable addressing various *RBM20* mutations by a single PE drug(*10, 12*). These features make *RBM20* an appealing, proof-of-concept target for *in vivo* cardiac PE.

In this study, we engineered and optimized *in vivo* PE for cardiomyocyte editing post systemic administration. When tested at the mouse *Rbm20* locus, our PE therapeutic resulted in efficient editing in cardiomyocytes and rescued cardiomyopathy phenotypes in the heterozygous *Rbm20^R636Q^*(corresponding to human *RBM20^R634Q^* mutation) DCM mouse model(*3*), representing, to our knowledge, the first demonstration of *in vivo* PE efficacy in cardiac indications. Additionally, we report a novel humanized mouse model carrying human WT or R634Q mutant sequences in and flanking the *RBM20* mutational hotspot region. This humanized model exhibits *RBM20* cardiomyopathy phenotypes and enables *in vivo* testing of PE targeting human *RBM20*. Finally, we demonstrated that our human *RBM20* PE therapeutic efficiently corrected the pathogenic mutation in the heart, rescued phenotypes in our humanized *RBM20* mouse model, and exhibited no detectable editing in the liver. Our study establishes an *in vivo* cardiac PE platform and exemplifies its therapeutic potential.

## RESULTS

### A dual-AAV based strategy is established and enables *in vivo* cardiac PE

To express prime editor protein *in vivo* in cardiomyocytes, we employed a 400bp human *TNNT2*-derived promoter which has been shown to mediate strong and cardiac selective gene expression(*23*). We chose the 1024-CFP intein-split PEmax DRNase-H design (split-PE) for dual-AAV based delivery(*15*). While Davis et al. arranged the pegRNA and the ngRNA transcription units both in the second, C-terminal AAV cassette (PE-C) and used human U6 and mouse U6 promoters for pegRNA and ngRNA, respectively, to avoid long stretches of homology on a single AAV genome that can lead to recombination(*15*), the compact size of our *TNNT2*-derived promoter enabled inclusion of the ngRNA transcription unit in the first, N-terminal AAV cassette (PE-N) and use of human U6 promoter for both guide RNAs (Fig. 1A).

**Fig. 1.**
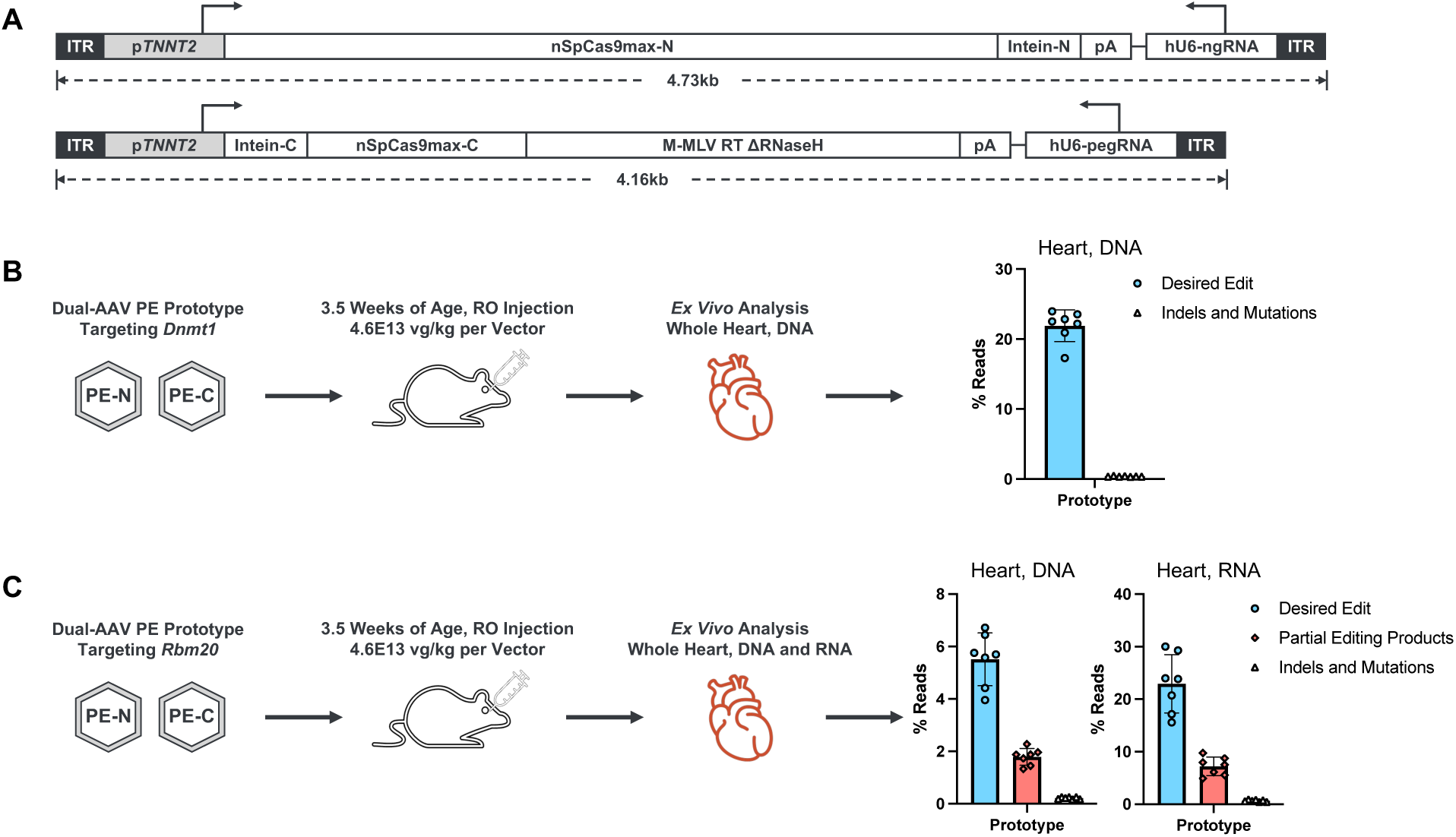
Establishing a dual-AAV based strategy for *in vivo* cardiac PE. (**A**) Schematic of our prototypical design of dual-AAV based PE cassettes. (**B**) *In vivo* cardiac editing at *Dnmt1* locus by the prototype. (**C**) *In vivo* cardiac editing at *Rbm20* locus by the prototype. PE-N and PE-C cassettes were packaged in TNC734 capsid. Seven mice were enrolled per group. *Ex vivo* editing analysis was performed at 17 weeks post-injection. Dots represent individual mice and error bars represent mean ± SD.

To assess *in vivo* cardiac editing efficiency, we chose a previously published pegRNA/ngRNA pair targeting the mouse *Dnmt1* locus(*15*) (Table S1) and packaged the PE cassettes in TNC734 capsid, an in-house engineered capsid with enhanced *in vivo* cardiomyocyte tropism in mouse following systemic administration (Fig. S1, A and B). Administered at 4.6E13 vg/kg per vector in 3.5-week-old mice via retro-orbital (RO) injection, our prototype PE editor showed 21.9% edited next-generation sequencing (NGS) reads at the *Dnmt1* locus in the heart, providing proof of concept for dual AAV vector-based *in vivo* cardiac PE (Fig. 1B).

We then applied our prototype to a disease-relevant locus, the corresponding human mutational hotspot in mouse *Rbm20*, and mimicked the real word application by designing a longer reverse transcription template in pegRNA that has the potential to cover the whole hotspot and to correct various pathogenic mutations (“mRbm20-PE prototype”). Carrying in-house optimized guide RNAs (Table S1), mRbm20-PE prototype functions *in vitro* but showed a lower editing efficiency compared to PE at the *Dnmt1* locus (Fig. S1, C to E). Consistent with *in vitro* results, our mRbm20-PE prototype edited the mouse *Rbm20* locus *in vivo* with 5.51% of reads carrying the desired edit when assessed from heart DNA (Fig. 1C and Fig. S1F), less efficiently than PE at the *Dnmt1* locus. It is worth noting that the expression of our prime editor is cardiomyocyte-selective due to the use of *TNNT2*-derived promoter, but genome copies from all cardiac cell types (including cells in which prime editor is not expressed) were collected and contributed to the denominator in our DNA-based analysis, which can cause underestimation of editing efficiency(*3, 4*). *Rbm20* is also primarily expressed in cardiomyocytes and *Rbm20* RNA transcripts recovered from whole heart lysate mainly originate from cardiomyocytes, so we reasoned that measuring *Rbm20* editing efficiency at the RNA level should better reflect the actual efficiency of our cardiomyocyte-selective PE design(*3, 4*). Our mRbm20-PE prototype yielded 22.9% desired editing in *Rbm20* RNA transcript analysis (Fig. 1C and Fig. S1G). In this (Fig. S1H) and other experiments (reported later in this manuscript), editing efficiency measured at the RNA level was consistently 4- to 5-fold of the efficiency measured at the DNA level for a cardiomyocyte-specific gene locus, which implies that around 20% to 25% of all genome copies recovered in our whole heart DNA extraction originated from cardiomyocytes. Thus, our prototype reaches near maximal cardiomyocyte editing at *Dnmt1* locus at this condition (21.9% edited in DNA-based analysis, Fig. 1B); however, our *in vivo* cardiac PE still needed optimization to enhance editing in more challenging settings, such as the *Rbm20* case.

To improve PE efficiency, we fused the RNA-binding, N-terminal domain of the small RNA-binding exonuclease protection factor La to the prime editing protein machinery(*24*). The resulting design, V19, functions *in vitro* and still allows for dual-AAV packaging (Fig. 2A and Fig. S1, C, D, E, I, and J). To avoid maxing out *in vivo* cardiac editing efficiency and allow head-to-head comparison of the prototype and V19 at the *Dnmt1* locus, we lowered the dosage to 1E13 vg/kg per vector and observed higher editing efficiency by V19 (17.5%) compared to the prototype (11.8%) in *Dnmt1* heart DNA analysis (Fig. 2B).

**Fig. 2.**
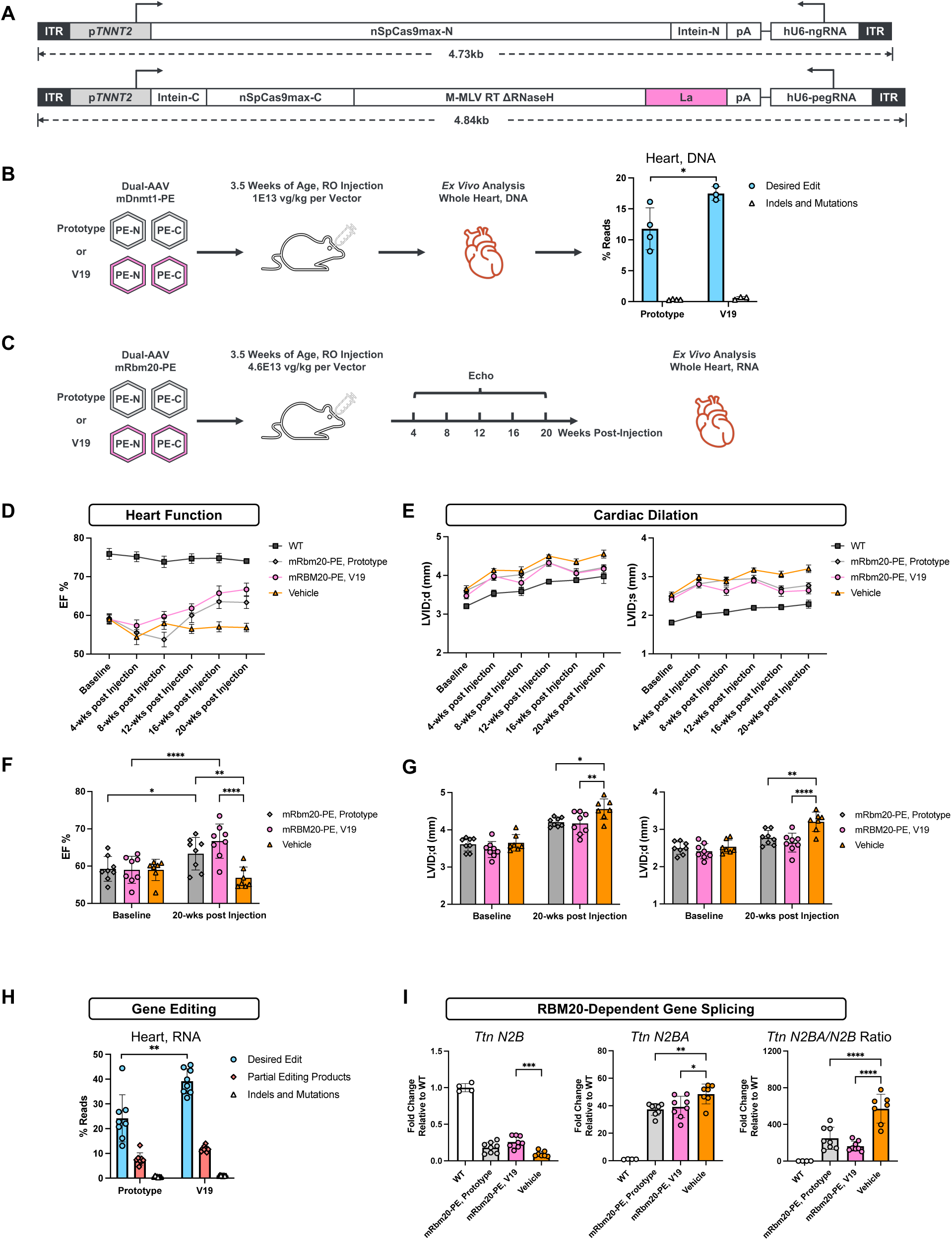
V19 mediates greater *in vivo* cardiac editing efficiency than the prototype and rescues phenotypes in *Rbm20^R636Q^* mouse model. (**A**) Schematic of V19 design. (**B**) *In vivo* cardiac editing at the *Dnmt1* locus by the prototype and V19. PE cassettes were packaged in TNC734 capsid. Three to four mice were enrolled per group. *Ex vivo* editing analysis was performed at 3 weeks post-injection. (**C**) Schematic design of *in vivo* editing efficiency and efficacy study testing mRbm20-PE prototype and V19 in *Rbm20^R636Q^* mouse model. PE cassettes were packaged in TNC755 capsid. Seven to eight mice were enrolled per group. (**D** to **G**) Ejection fraction (EF) measurements (D and F) by Echo and left ventricle internal diameter measurements (E and G) at diastolic stage (LVID;d) and at systolic stage (LVID;s). For D and E, error bars represent mean ± SEM. (**H**) Editing efficiency at the *Rbm20* locus, detected from whole heart RNA samples. (**I**) Splicing patterns of *Ttn* gene. For B and H, dots represent individual mice, error bars represent mean ± SD, statistical significance was calculated using unpaired t-test. For E, G, and I, dots represent individual mice, error bars represent mean ± SD, statistical significance was calculated using two-way ANOVA with Tukey’s multiple comparisons test. For B and F to I, \**P* < 0.05, \*\**P* < 0.01, \*\*\**P* < 0.001, \*\*\*\**P* < 0.0001.

### *In vivo* cardiac PE improves heart function of mouse *Rbm20^R636Q^* DCM model

We next performed an *in vivo* editing efficiency and efficacy study in *Rbm20^R636Q/+^* mice, an established model of human *RBM20* cardiomyopathy corresponding to human *RBM20^R634Q^*mutation(*3*), to compare mRbm20-PE V19 against mRbm20-PE prototype and assess the therapeutic potential of *in vivo* cardiac PE (Fig. 2C). We used TNC755 capsid (Fig. S1, A and B) in this and all following *in vivo* experiments described in the manuscript. Prior to RO administration at 4.6E13 vg/kg per vector dosage at 3.5 weeks of age, baseline echocardiographic (Echo) analyses were performed in *Rbm20^R636Q/+^* mice and in age-matched wildtype (WT) controls, and their cardiac function was subsequently monitored by Echo at 4-week intervals up to 20 weeks post-injection, followed by *ex vivo* analyses (Fig. 2C).

Vehicle treated *Rbm20^R636Q/+^* mice exhibited reduced ejection fraction and enlarged left ventricle internal diameters compared to WT mice (Fig. 2, D and E), consistent with the DCM phenotypes reported by Nishiyama et al(*3*). Both mRbm20-PE prototype and mRbm20-PE V19 treatments improved ejection fraction (Fig. 2, D and F) and mitigated cardiac dilation (Fig. 2, E and G) over time. The rescue of heart function and dilation trended better in mRbm20-PE V19 treated mice compared to mRbm20-PE prototype treated mice (Fig. 2, D to G). Consistently, *ex vivo* editing efficiency analysis detected on average 39.1% and 24.2% of mouse *Rbm20* transcripts carrying the desired edit in the hearts of mRbm20-PE V19 and mRbm20-PE prototype treated mice, respectively (Fig. 2H and Fig. S2, A to C). *Ex vivo* splicing analysis of RBM20 downstream targets Titin (*Ttn*) and calcium/calmodulin dependent protein kinase II delta (*Camk2d*) revealed partial rescue of abnormal splicing patterns(*3, 25*) by mRbm20-PE V19 and mRbm20-PE prototype treatment (Fig. 2I and Fig. S2D).

The heterozygous genotype of *Rbm20^R636Q/+^* mice complicates the interpretation of editing efficiency as both *Rbm20^R636Q^*and *Rbm20* WT alleles can be recognized and edited to the same final sequence by mRbm20-PE, and we cannot directly measure the percentage of mutant allele copies corrected in NGS analysis. The relative abundance of unedited mutant and WT allelic sequences stays roughly unchanged (close to 1:1 ratio) in our mRbm20-PE studies regardless of whether PE was performed (Fig. S1G and Fig. S2A), suggesting that our mRbm20-PE editors edit *Rbm20^R636Q^* and *Rbm20* WT alleles at similar efficiencies and the overall percentage of desired edit observed in our NGS analysis is a close approximation of the percentage of mutant allele copies corrected.

Together, these results demonstrate the improvement of V19 design over the prototype and support the therapeutic potential of *in vivo* cardiac PE; however, unedited *Rbm20^R636Q^* allelic sequence still accounted for 33.8% of all reads in NGS analysis of mRbm20-PE V19 treated mice, which translates to nearly 70% of cardiomyocytes still carrying the uncorrected mutant allele, prompting us to further optimize the *in vivo* PE design.

### Further optimization improves *in vivo* cardiac PE efficiency

While Yan et al. showed that fusing the RNA-binding domain of La to the C-terminus of PE protein, namely PE7, was the most efficient among all fusing locations tested(*24*), we found that fusing the RNA-binding domain internally between Cas9 and M-MLV RT DRNase-H showed improved efficiency over V19, which has the C-terminal fusion arrangement, under our split-PE setting. We name this new design V20 (Fig. 3A and Fig. S1, C, D, E, I, and J). When tested *in vivo* at 2.8E13 vg/kg per vector, V20 more than doubled cardiac editing efficiency compared to V19 at the mouse *Rbm20* locus (74.5% versus 34.5%, measured at RNA transcript level) (Fig. 3B and Fig. S3A).

**Fig. 3.**
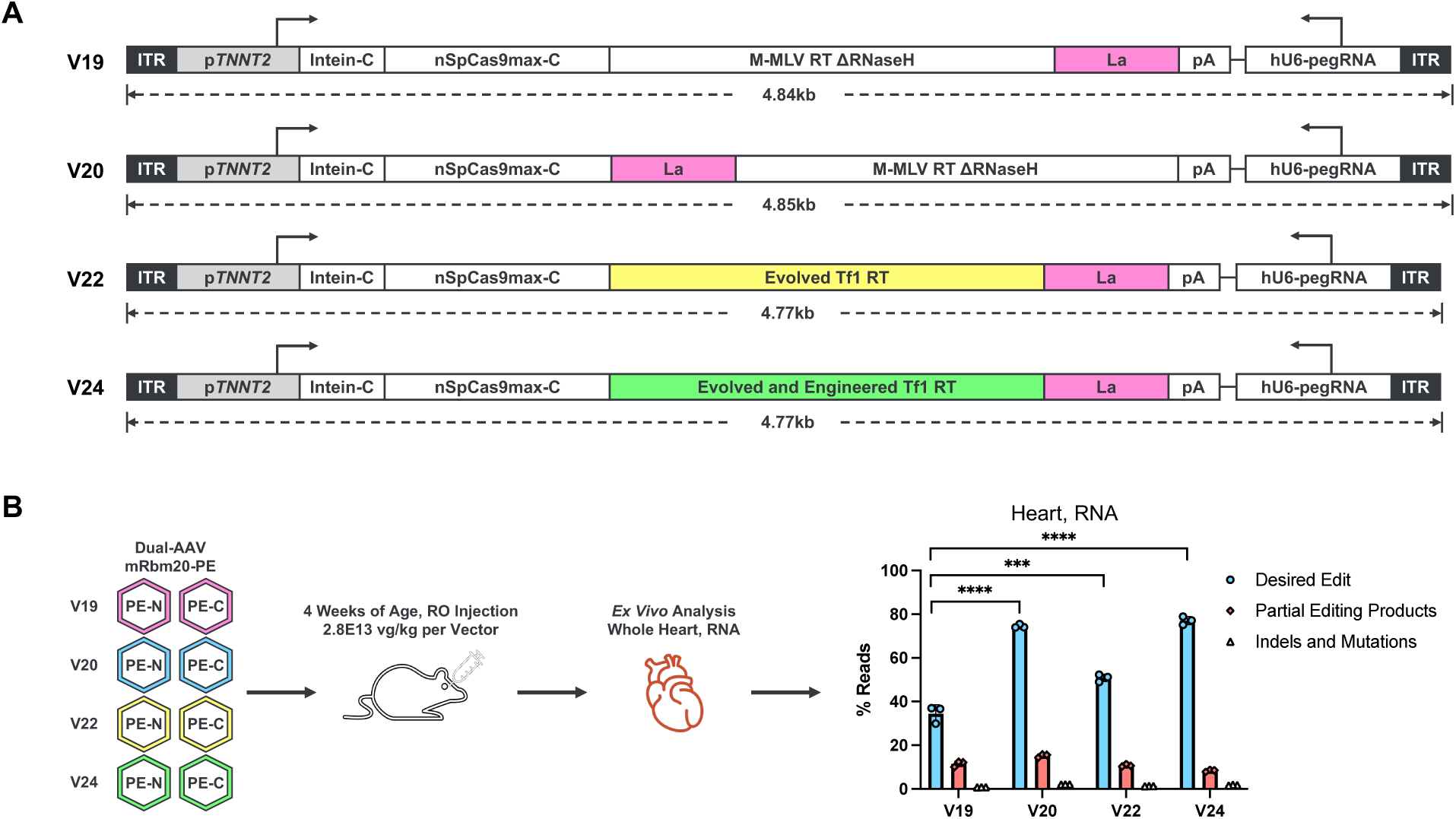
Further optimization enhances *in vivo* cardiac editing efficiency. (**A**) Schematics of V19, V20, V22, and V24 designs. (**B**) *In vivo* cardiac editing at the *Rbm20* locus by mRBM20-PE V19, V20, V22, and V24. WT C57BL/6J mice were used in this study. *Ex vivo* analysis was performed at 8 weeks post-injection. Dots represent individual mice, error bars represent mean ± SD, statistical significance was calculated using one-way ANOVA with Tukey’s multiple comparisons test. \*\*\**P* < 0.001, \*\*\*\**P* < 0.0001.

In parallel, we explored alternative reverse transcriptases developed by Doman et al(*26*). By swapping M-MLV RT DRNase-H in V19 to evolved Tf1 RT and evolved and engineered Tf1 RT (used in PE6b and PE6c, respectively, in Doman et al.), we developed V22 and V24, respectively (Fig. 3A and Fig. S1, C, D, E, I, and J). While V22 yielded 50.8% of mRbm20 RNA transcripts carrying the desired edit *in vivo* in mouse heart, V24 showed 77.1% editing, comparable to V20 and more than doubled compared to V19 (Fig. 3B and Fig. S3A).

Furthermore, we combined the internal fusion arrangement of La of V20 with the alternative reverse transcriptases in V22 and V24. The resulting PE designs, V27 and V28, functioned well *in vitro* (Fig. S1, C, D, E, I, and J).

Finally, long-term exposure to PE activity may not always be beneficial and in certain cases, inactivation of PE expression and/or function post-administration may be desirable. To this end, we propose a strategy of PE self-inactivation (SI) (Fig. S3B) and provide an *in vivo* proof-of-concept demonstrating editing at both the genomic target and the SI site (Fig. S3, C and D).

### A novel humanized *RBM20* mouse model displays *RBM20* cardiomyopathy phenotypes

Having demonstrated the therapeutic potential of *in vivo* cardiac PE and having identified PE designs with enhanced *in vivo* efficiency, we sought to apply this platform to treating human pathogenic mutations. However, the differences at the nucleotide sequence level of orthologous genes in different species often impede testing of human gene targeting PE in animal models. We overcame this challenge by designing and generating a novel humanized *RBM20* mouse model.

The *RBM20* mutational hotspot region is located near the beginning of exon 9. Taking advantage of the high conservation between human RBM20 and mouse Rbm20 protein sequences in and flanking the mutational hotspot region, we replaced 64 base pairs (bp) at the 5’ start of exon 9 of mouse *Rbm20* with the corresponding sequence of human *RBM20* (Fig. 4 and Fig. S4A). Additionally, we humanized the last 200 bp of intron 8, making the total humanized region to 264-bp-long, which allows binding by pegRNAs and ngRNAs designed for human *RBM20* editing. To enable disease modeling and *in vivo* efficacy testing, we constructed two lines carrying humanized *RBM20* WT sequence or humanized *RBM20^R634Q^* sequence (human RBM20 amino acid position numbering is used here to avoid confusion, despite that the actual mutation position is 636 in the human-mouse chimeric gene) (Fig. 4 and Fig. S4A), and further generated heterozygous *hRBM20^R634Q^/hRBM20^+^*mice by crossing these two humanized lines, confirmed by genotyping and Sanger sequencing (Fig. S4B).

**Fig. 4.**
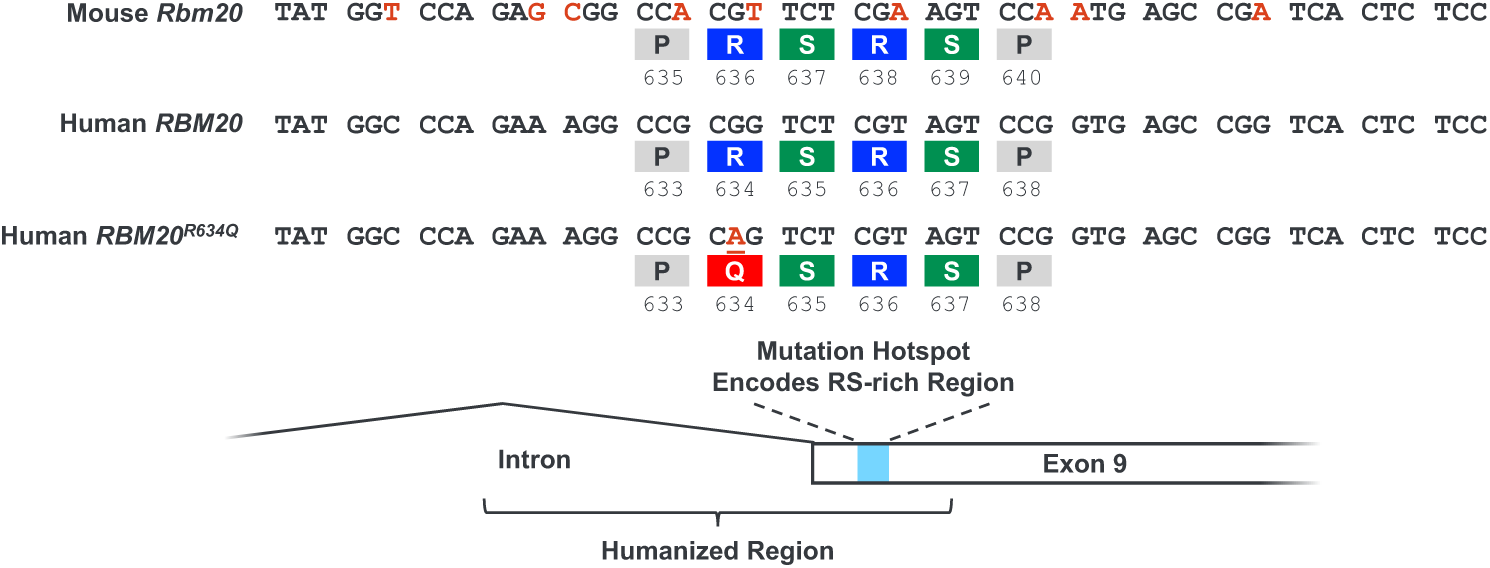
Design and generation of a partially humanized *RBM20* mouse model. Schematic of the design of our humanized *RBM20* mouse model, highlighting the sequence differences between mouse and human, and the location of the humanized region.

To characterize this newly generated mouse model, we conducted comprehensive phenotypic assessments. Echo revealed that while humanized WT controls remained healthy, heterozygous mutant mice (*hRBM20^R634Q^/hRBM20^+^*) developed a moderate DCM phenotype with reduced ejection fraction (EF) and increased left ventricular internal diameter in systole (LVID;s) and diastole (LVID;d) (Fig. 5, A and B). Homozygous mutants (*hRBM20^R634Q^/hRBM20^R634Q^)* exhibited severe heart failure and ventricular enlargement (Fig. 5, A and B). Notably, the homozygous mice showed significantly increased heart weight to body weight (HW/BW) ratios (Fig. 5C) and a shortened lifespan (<3 months) (Fig. 5D). Visualization of freshly excised cardiac tissue and H&E-stained heart slices revealed dilation in 22-week-old heterozygous mutant mice (Fig. 5, E and F). Masson’s trichrome staining showed no notable fibrosis (Fig. 5G). Molecular analysis confirmed abnormal splicing of *Ttn* (increased N2BA isoform, decreased N2B isoform; Fig. 5H) and *Camk2d* (increased Camk2d-A, decreased B/C isoforms; Fig. 5I) in mutant hearts. Additionally, the expression levels of heart failure markers natriuretic peptide A and B (*Nppa* and *Nppb*), as well as the fibrosis-associated gene transforming growth factor, beta (*Tgf-β*), were modestly increased in heterozygous hearts (Fig. S5, A and B). Electrocardiogram (EKG) analysis did not reveal any significant differences between mutant and wild-type mice (Fig. S5C). Overall, our novel humanized *RBM20* mouse model exhibits *RBM20* cardiomyopathy phenotypes reminiscent of those observed in mouse endogenous *Rbm20* disease model and human patients(*3, 27*), and provides a valuable platform for evaluating the functional consequences of human *RBM20* mutation in a physiological context and for testing PE targeting human *RBM20*.

**Fig. 5.**
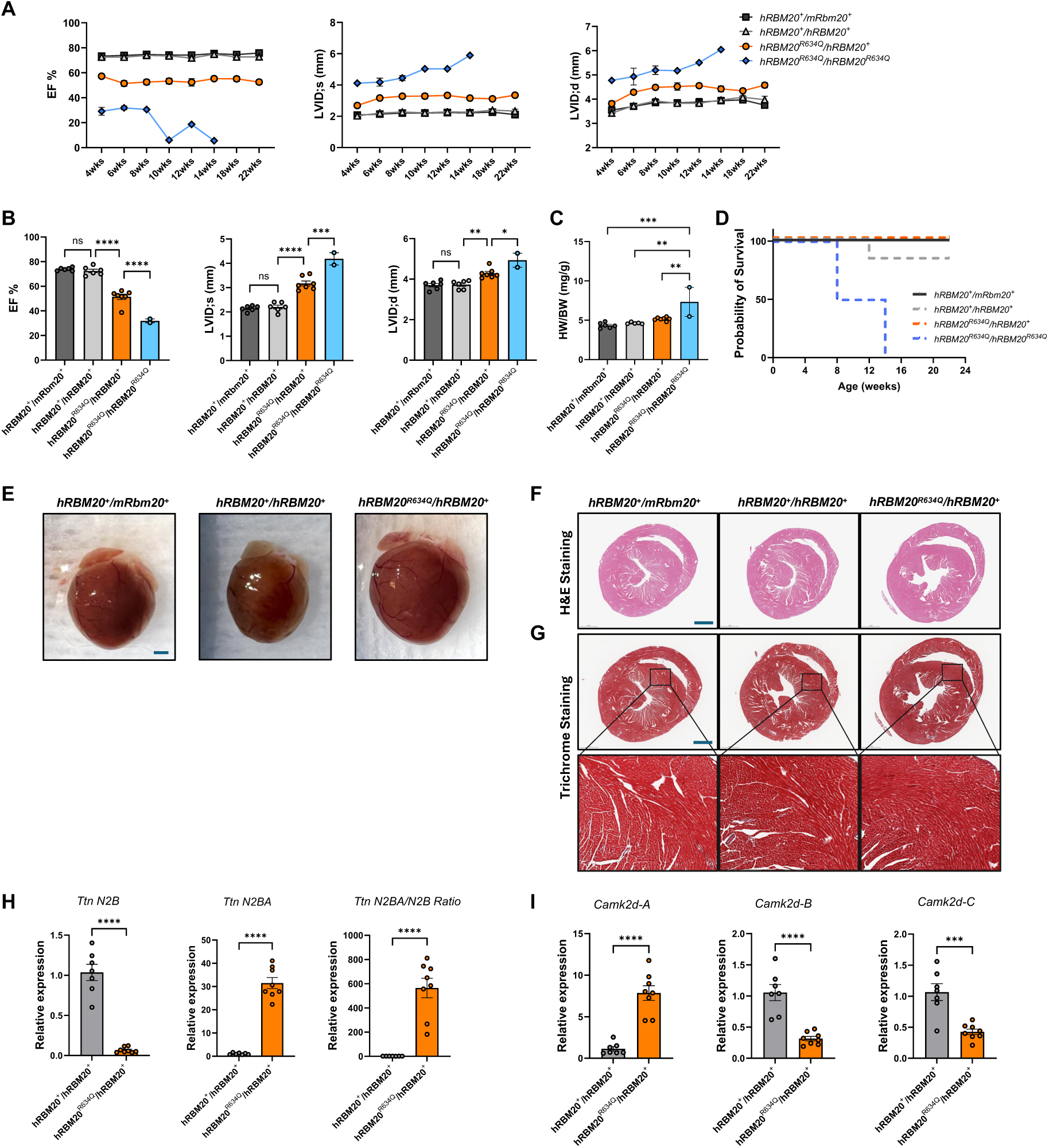
Humanized *RBM20^R634Q^*mice show DCM phenotype. (**A**) Cardiac function progression of heterozygous humanized WT *(hRBM20^+^/mRbm20^+^)* (n=6), homozygous humanized WT *(hRBM20^+^/hRBM20^+^)* (n=6), heterozygous mutant *(hRBM20^R634Q^/hRBM20^+^)* (n=7), and homozygous mutant *(hRBM20^R634Q^/hRBM20^R634Q^)* (n=2). (**B**) Ejection fraction (EF), left ventricle internal diameter at systolic stage (LVID;s) and left ventricular internal diameter at diastolic stage (LVID;d) at 6 weeks of age. Heart weight to body weight ratios (HW/BW) at death (animals died during the study) or at 22 weeks of age. For **b** and **c**, dots represent individual mice. For A to C, error bars indicate mean ± SEM. and statistical significance was calculated using one-way ANOVA with Tukey’s multiple comparisons test. (C) Mortality over time. (**E**) Representative whole-heart images. (**F**) Representative hematoxylin and eosin (H&E)–stained heart sections. (**G**) Representative Masson’s trichrome–stained heart sections (n = 5–7 per group). For E to G, scalebar = 1 mm. (**H**) Splicing patterns of *Ttn*. (**I**) Splicing patterns of *Camk2d*. For H and I, dots represent individual mice, error bars indicate mean ± SEM., and statistical significance was calculated using unpaired t-test. For B, H, I, \**P* < 0.05, \*\**P* < 0.01, \*\*\**P* < 0.001, \*\*\*\**P* < 0.0001.

### *In vivo* cardiac PE corrects human *RBM20^R634Q^* mutation and rescues cardiomyopathy phenotypes

To evaluate *in vivo* PE of human *RBM20*, we administered PE-hRBM20-4.9 (Table S2), which uses V20 design and carries a pair of human *RBM20*-targeting pegRNA/ngRNA (Table S1) developed and validated *in vitro* (Fig. S1I), to *hRBM20^R634Q^/mRbm20^+^*mice at 1.5E13, 3E13, and 5E13 vg/kg dosages. *Ex vivo* editing analysis at 4 weeks post-injection detected 16.9%, 29.3%, and 32.5% *hRBM20* transcripts fully edited at these three dosages (Fig. S5, D to F), indicating diminishing returns on the editing dose response. PE-hRBM20-4.9 installs a silent edit in addition to the mutation-correcting edit (Fig. S5D) and the design of our mRbm20-PE also followed the same concept (Fig. S1F). This allows us to validate the PE constructs *in vitro* in WT human cells prior to *in vivo* experiments, however, in both cases we observed large amounts of partial editing products with only the silent edit but not the mutation-correcting edit (Fig. 1C, Fig. 2G, Fig. S1F, and Fig. S5, D to F). The proximity to the nick site may make the silent edit more likely to be installed than the mutation-correcting edit (for example, due to premature stop of reverse transcription), and the overlapping of the silent edit with the protospacer adjacent motif (PAM) or spacer binding region may leave silent-edit-only DNA harder to be recognized again and acquire the mutation-correcting edit. Thus, we developed two new *RBM20*-targeting pegRNAs (Table S1) which edit mutant allele to WT sequence without incorporating any silent edit. We validated their performance in mouse embryonic fibroblasts (MEFs) isolated from homozygous *hRBM20^R634Q^/hRBM20^R634Q^*mice and named the resulting PE constructs PE-hRBM20-4.11 and PE-hRBM20-4.12 (Fig. S5G and Table S2).

Encouraged by the performance of our human *RBM20* targeting PE constructs and the clear *RBM20* cardiomyopathy phenotypes of our novel humanized mouse model, we performed an efficacy study in which humanized heterozygous *hRBM20^R634Q^/hRBM20^+^* mutant mice were administered systemically with PE-hRBM20-4.9, PE-hRBM20-4.11, or PE-hRBM20-4.12 at 3E13 vg/kg at 3.5 weeks of age. Heart function and dilation were monitored by echocardiography prior to dosing and at 4-week intervals for up to 16 weeks post-injection, followed by *ex vivo* editing and gene splicing analyses (Fig. 6A). While vehicle treated *hRBM20^R634Q^/hRBM20^+^*mice displayed reduced ejection fraction and increased left ventricle internal diameters, consistent with model development observations, PE-hRBM20-4.9, PE-hRBM20-4.11, and PE-hRBM20-4.12 treated animals showed better heart function and cardiac dilation compared to the vehicle control (Fig. 6, B to E). PE-hRBM20-4.11 showed the strongest rescue among treated groups, with both ejection fraction and cardiac dilation measurements improved over time trending towards the WT control (*hRBM20^+^/hRBM20^+^*) levels (Fig. 6, B to E). Since the desired post-editing product of PE-hRBM20-4.11 and PE-hRBM20-4.12 is identical to and indistinguishable from the pre-existing WT hRBM20 allelic sequence (Fig. S6, A to C), we report the percentages of reads that encode WT hRBM20 protein or R634Q mutant protein in NGS editing analysis of this study. While nearly half (49.0%) of all reads in vehicle treated heterozygous *hRBM20^R634Q^/hRBM20^+^*mice corresponded to R634Q mutant, PE-hRBM20-4.9, PE-hRBM20-4.11, and PE-hRBM20-4.12 treated groups had on average 27.9%, 16.4%, and 17.9% reads, respectively, encoding R634Q mutant protein (Fig. 6, F and G, and Fig. S6C), which can be interpreted as roughly 57%, 33%, and 37% of all cardiomyocytes with the mutant allele remaining unedited. The splicing patterns of RBM20 downstream targets *Ttn* and *Camk2d* were also partially rescued in all three treated groups, with the PE-hRBM20-4.11 trending the best (Fig. 6, H and I).

**Fig. 6.**
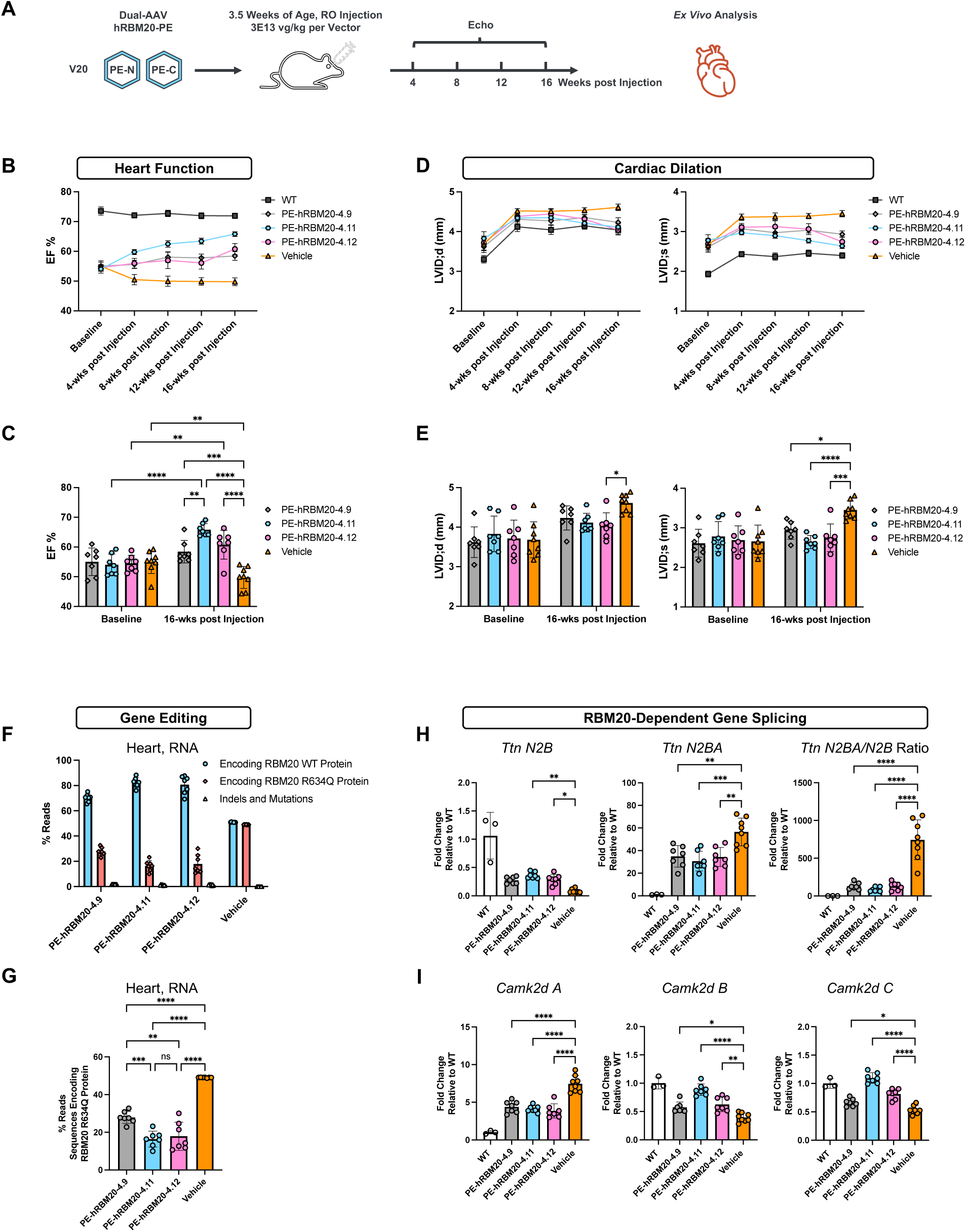
PE therapeutic targeting human *RBM20* mediates *in vivo* correction of pathogenic mutation in the heart and rescues cardiomyopathy phenotypes in humanized *RBM20^R634Q^*mouse model. (**A**) Schematic design of *in vivo* editing efficiency and efficacy study testing PE-hRBM20-4.9, PE-hRBM20-4.11, and PE-hRBM20-4.12 in our novel humanized mouse model. PE cassettes were packaged in TNC755 capsid. Seven to eight mice were enrolled per group. (**B** and **C**) Ejection fraction (EF) measurements by Echo. (**D** and **E**) Left ventricle internal diameter at diastolic stage (LVID;d) and at systolic stage (LVID;s) measurements. (**F** and **G**) Editing efficiency at humanized *RBM20* locus, detected from whole heart RNA samples. (**H**) Splicing patterns of *Ttn*. (**I**) Splicing patterns of *Camk2d*. For B and D, error bars represent mean ± SEM. For C to I, dots represent individual mice, error bars represent mean ± SD. For C and E, statistical significance was calculated using two-way ANOVA with Tukey’s multiple comparisons test. For G to I, statistical significance was calculated using one-way ANOVA with Tukey’s multiple comparisons test. For C, E, G to I, \**P* < 0.05, \*\**P* < 0.01, \*\*\**P* < 0.001, \*\*\*\**P* < 0.0001.

Systemic administration of dual-AAV prime editors expressed by ubiquitous promoters leads to high levels of editing in the liver(*15, 17*). To assess whether our *in vivo* cardiac PE therapeutic shows off-targeting to the liver, we extracted DNA from the liver of PE-hRBM20-4.9, PE-hRBM20-4.11, PE-hRBM20-4.12, and vehicle treated animals and performed editing efficiency analysis. The editing was undetectable at the target locus in the liver of all three treated groups, in contrast to what was observed in heart DNA samples (Fig. S6, D to G). We next investigated deeper into PE-hRBM20-4.9 treated samples in which we can clearly distinguish several post-editing sequences from the pre-existing WT allele thanks to the silent edit introduced by PE-hRBM20-4.9. We saw no detectable change in any post-editing sequence populations (Fig. S6H).

Finally, we observed very low levels of indels and undesired mutations in this (1% to 2% of heart RNA reads and 0.2% to 0.5% of heart DNA reads) (Fig. 6F and Fig. S6, C and D) and all previous *in vivo* PE experiments, which is consistent with the observation by Davis et al(*15*).

Together, these results demonstrate that our *in vivo* cardiac PE corrects the pathogenic mutation in the heart and rescues cardiomyopathy phenotypes in our novel humanized *RBM20* mouse model with no detected editing in the liver.

## DISCUSSION

We present a dual-AAV based *in vivo* cardiac PE platform with designs and optimizations for targeted and efficient *in vivo* PE in cardiomyocytes. Our results suggest that while incorporating the RNA-binding domain of La enhances editing efficiency, the location of La in the fusion protein has a large impact on *in vivo* cardiac PE efficiency, at least for the genomic target locus tested in this study. Replacing MLV RT with Tf1 RT variants also improves editing efficiency, and interestingly, lowered the tendency to generate partially edited products, compared to their counterpart using MLV RT in the mouse *Rbm20* case study where PE installs more than one nucleotide change (Fig. S3A). Future study may investigate whether this trend is consistent across different targets and explore the underlying mechanisms. Additionally, the promising *in vitro* results of our V27 and V28 designs, which combine the favorable location of La and Tf1 RT variants, warrant future *in vivo* testing.

For split-PE to function, both PE-N and PE-C need to be present in the same cell and expressed at a sufficiently high level, posing a high demand for delivery efficiency. Consistently, our in-house engineered cardiac tropic capsids mediated enhanced PE efficiency compared to the same PE design delivered by AAV9, especially at a low dose level (Fig. S1B). Moreover, the efforts of identifying compact nucleases and reverse transcriptases(*26, 28–30*) may enable single-AAV PE in the future, potentially improving efficacy and/or reducing dosage.

Although we developed our *in vivo* PE platform aiming at cardiac editing, our PE designs can be adopted for targeting other organs and tissues. Researchers may modify our PE designs by changing the promoter and/or the capsid to those that better fit their organs/tissues of interest. Thus, our PE optimization efforts possess broader impact beyond cardiac indications.

Here we use *RBM20^R634Q^* mutation as a case study, however, our hRBM20-PE therapeutic was designed to be compatible with various mutations in or near the *RBM20* mutational hotspot region, including but not limited to P633L, R634W, R634Q, S635A, R636C, R636S, R636H, S637G(*21*), offering an attractive advantage over BE strategies(*3, 4*). Future *in vitro* and *in vivo* studies should be performed to investigate the editing efficiency of our hRBM20-PE therapeutic on other *RBM20* mutations.

In our humanized *RBM20* efficacy study, the heart function and left ventricle dilation were still trending better in two of the treatment groups at the end of in-life observation stage (Fig. 6, B and D). AAV delivery supports long-term transgene expression(*31*), thus the percentage of unedited mutant *RBM20* copies may continue to decrease over time, which would lead to continuous improvement of efficacy. It is also possible that the phenotypic improvements lag after the correction of mutant *RBM20* copies. Therefore, future editing kinetics investigation may be of interest.

Finally, while we employed our novel humanized *RBM20* mouse model for *in vivo* PE testing, this model also enables *in vivo* testing of other sequence-specific therapeutics designed to target the human *RBM20* mutational hotspot, including but not limited to CRISPR/Cas9 editing, BE, antisense oligonucleotides, RNA interference, and CRISPR interference. Additionally, it may serve as a valuable DCM model to test various small molecule and gene replacement drugs.

There are some limitations to this work, raising the need of future experiments for our findings to be applied in the clinic. First, we performed *in vivo* editing and efficacy studies in mice and it remains unclear whether our *in vivo* cardiac PE platform can achieve efficient editing in large animals. Investigations in non-human primate models should be pursued. Second, long-term follow-up studies addressing the endurance of efficacy, immune responses, off-targeting, tumorigenesis, etc., were not included in the current research but should be performed prior to clinical translation. In addition, only one dosage was tested in the *RBM20 in vivo* efficacy experiment. It would be a crucial future step to examine the dose response and tolerance of *in vivo* cardiac PE therapeutic. Last, we focused on the technological development of *in vivo* cardiac PE and the demonstration of cardiac function improvements in the current study, whereas biological and mechanistic questions, such as the intracellular localization of RBM20 protein and whole transcriptome features pre- and post-treatment, may be of interest in future studies.

## MATERIALS AND METHODS

### Study design

This study aimed to develop an *in vivo* cardiac PE platform and test its therapeutic potential. This study used *RBM20* cardiomyopathy as a case study to demonstrate the *in vivo* efficacy of our PE platform. Post genotyping, animals were randomized and assigned to experimental groups, aiming at having similar average baseline ejection fraction values and body weight across groups. In efficacy experiments, injection and echocardiography analysis were performed by blinded operators. Each experiment carried biological replicates as indicated in the figures and figure legends.

### Molecular cloning

Sequences of guide RNAs, plasmids, AAV transgene cassettes, and AAV capsids are available upon request.

Plasmids were generated via Gibson assembly or ligation, using NEBuilder HiFi DNA Assembly (NEB E2621) or Quick Ligase (NEB M2200S). Restriction digests of backbone plasmids were assembled with or ligated to GenScript gene fragments or polymerase chain reaction (PCR) amplicons. Plasmids were purified from Stellar Competent *Escherichia coli* (Takara 636763) using Azenta’s standard plasmid prep service or using PureYield Plasmid Miniprep kit (Promega A1223).

### Cell culture

HEK293T cells (American Type Culture Collection (ATCC), CRL-3216), Neuro-2A cells (ATCC, CCL-131), and MEF cells were grown in DMEM plus GlutaMAX (Gibco 10566-016) supplemented with 10% (v/v) FBS (Gibco A56707-1) and 5% (v/v) Anti-Anti (Gibco 15240-062) at 37°C with 5% CO_2_ unless otherwise stated. MEF cells were isolated from humanized *RBM20^R634Q^* mutant homozygous mouse embryo harvested at E13-E15. Embryo was covered in DPBS (Gibco 14190-144), and placenta and maternal tissue were removed. Top of head was cut away and innards removed. Head was retained for DNA isolation for genotyping. Embryo body was placed in 10cm plate (Corning 430167), 5mL 5x trypsin was added, and embryo was minced with razor blade and incubated at 37°C for 30-45 min. Trypsin activity was quenched with 5mL MEF media (DMEM with 10% (v/v) FBS and 5% (v/v) Pen Strep (Gibco 15140-122)), and tissue was broken up by pipetting 10-20x. Sample was centrifuged, resuspended in 10mL MEF media, plated in 10cm plate, and allowed to grow to confluency (3-4 days).

### Transfections of HEK293T, Neuro-2a, and MEF Cells

Twenty-four hours before transfection, HEK293T and Neuro-2a cells at more than 90% viability were seeded on 96-well plates (Thermo Fisher Scientific FB012931) at a density of 10,000-12,000 cells per well. HEK293Ts were transfected with 0.3 μl of Lipofectamine 3000 (Thermo Fisher Scientific L3000001) and 0.2 μl of P3000 with 100 ng of DNA in DMEM plus GlutaMAX supplemented with 2% (v/v) FBS and 5% (v/v) Anti-Anti. Neuro-2a cells were transfected with 0.35 μl Lipofectamine 2000 (Thermo Fisher Scientific 11668030) and 350 ng DNA. HEK293T and Neuro-2a cells were co-transfected with dual PE cassette plasmids and two linear DNA fragments encoding the ngRNA and pegRNA. Linear DNA fragments were amplified from plasmid or from GenScript gene fragments using Q5 Hot Start High-Fidelity 2X Master Mix (NEB M0494S) and primers [ACTCCATCACTAGGGGTTCCTG] and [TTGGGAAGACAATAGCAGGCATG]. PCR reactions were performed with the following conditions: 98°C 30 s; 12-18 cycles 98°C 10s, 67°C 20s, 72°C 20 sec; 72°C 2 min and purified using QIAquick PCR Purification Kit (Qiagen 28106).

For HEK293T cell transfections, DNA was delivered in a 1:5 plasmid:linear DNA ratio, and for Neuro-2a cell transfections, DNA was delivered in a 1:10 plasmid:linear DNA ratio. For each replicate, 5 wells total were transfected (to be pooled after harvest) to average out potential well-by-well discrepancies in cell density. 72 hours post-transfection, media was removed and cells were harvested by adding 50 μl extraction buffer (Extraction Buffer (Abcam ab193970) + Extraction Enhancer (Abcam ab193971) + cOmplete Protease Inhibitor Cocktail (Roche 11697498001)) per well and pipetting up and down.

For MEF transfections, ∼80,000 cells per well were plated on 24-well plates (Corning 353047) 24 hours before transfection. MEFs were transfected at 80% confluency with 0.5 μl GenJet (SignaGen Laboratories SL100489-MEF), 500 ng DNA, and 0.5 μl CombiMag (Oz Biosciences CM20100) per well. DNA supplied was a single plasmid containing whole PE and both ngRNA and pegRNA. For each replicate, two wells total were transfected (to be pooled after harvest). After adding transfection mix, each plate was incubated on a magnetic plate (Oz Biosciences MF10000) for 20 min at 37°C. Wells were replaced with fresh media 24 hours post-transfection. 72 hours post-transfection, cells were detached using TrypLE Express (Gibco 12604-013), centrifuged at 300g for 5 min, and resuspended in 50 μl of extraction buffer.

For all cell lines, after harvest and pooling, samples were treated with proteinase K (Thermo Scientific EO0491) at 55°C for 16 h, followed by 95°C for 20 min. The resulting lysates served as gDNA templates in NGS editing efficiency analysis.

### gRNA selection

For selection of optimized pegRNAs and ngRNAs targeting mouse *Rbm20*, several rounds of screening were performed in Neuro-2a cells. For selection of optimized pegRNAs and ngRNAs targeting human *RBM20*, several rounds of screening were performed in HEK293T cells with validation of several tops hits in MEF cells from humanized *RBM20^R636Q^* mice. Variables modified for the pegRNA include nick-site, RTT and PBS lengths, spacer length, post-edit DNA sequence, and linker sequence. For the ngRNA, variables modified include nick-site and spacer length. Various combinations of pegRNAs and ngRNAs were tested. Transfections and harvesting were performed as described above.

### Recombinant AAV production

We follow the standard HEK293T triple-transfection rAAV production and iodixanol gradient ultracentrifugation purification protocol. Briefly, HEK293T cells were seeded to Corning HYPERFlask Cell Culture Vessels at 50M cells per HYPERFlask in DMEM (Gibco, 10566024) + 10% FBS (Gibco, 26140-095) + Antibiotic-Antimycotic (Gibco, 15240062). At ∼48 hours post seeding, 500 ug of DNA and 1 mg of PEI-MAX (Transporter 5 Transfection Reagent, Kyfora Bio, 26008-50) were mixed and transfected to the cells in DMEM + 2% FBS + Antibiotic-Antimycotic. For transfection DNA, 500 ug of pAdHelper + REP-CAP plasmid + transgene cassette plasmid (1:1:1 molar ratio) were used. At ∼72 hours post transfection, cells were harvested by mechanical shaking, pelleted by spinning, resuspended in 150mM NaCl, 50mM Tris-HCl, pH 8.5, and lysed by 4 freeze-thaw cycles. Next, non-encapsulated DNA in the lysate was removed by 1 hour of Benzonase (Millipore, E1014) digestion at 37°C. After a 20-minute centrifugation at 24,000 rcf, the supernatant was loaded to the top of an iodixanol gradient with 15%, 25%, 40%, and 60% layers. Following 1.5 hours of ultracentrifugation (Beckman Coulter Type 70 Ti rotor, 59,000 rpm), 40% layer near the 40%-60% interface was extracted by a 5mL sterile syringe attached to an 18-G needle. The extracted solution was desalted by Amicon® Ultra-15 100kDa centrifugal filter (Millipore, PFHYS1008) in HBSS + 0.001% Pluronic F68 (Gibco, 24040-032) and the purified rAAV viruses were titered by Quant-iT PicoGreen dsDNA assay (Invitrogen) following manufacturer’s instructions.

### Mouse studies

Animal studies were performed according to Tenaya Therapeutics’ animal use guidelines. The animal protocols were approved by the Institutional Animal Care and Use Committee (IACUC number: 2023.007). All mice enrolled had C57BL/6 genetic background. *Rbm20^R636Q^* knock-in mice were obtained from University of Texas Southwestern Medical Center, and the information on how they generated the mice was detailed in their publication. Cyagen Biomodels LLC (Santa Clara, CA) was contracted for generating humanized RBM20 WT and R634Q mutant knock-in mice utilizing CRISPR/Cas 9 system on C57BL/6N background. 2 male founders and 3 female founders were obtained for each line initially, humanized knock-in sequences in the F1 generation were confirmed by sequencing. Both male and female mice were enrolled in all studies. Ear tissue genomic DNA was extracted and used for genotyping.

### Echocardiography

Cardiac function was assessed by transthoracic echocardiography using high resolution micro-imaging systems (Vevo 3100 Preclinical Imaging System, VisualSonics). Briefly, anesthetized spontaneously breathing mice (1-3% isoflurane and 98.5-99% O2) were placed in the supine position on a temperature-controlled heating platform to maintain their body temperature at ∼37°C. Nair was used to remove hair and expose the skin to the probe. Parasternal short-axis M-mode tracings of left ventricle (LV) were recorded for LV ejection fraction (EF) and LV internal diameter in end systole and end diastole (LVID;s and LVID; d) calculation. Vevo Lab software was used for analyses.

### Electrocardiogram recordings

Mice were anesthetized with 1-1.5% isoflurane and 98.5-99% O2 via a nose cone (following induction in a chamber containing 3-4% isoflurane in oxygen). Body temperature was monitored continuously and maintained at 37–38 °C using a heat pad. Two lead ECG (leads II) were recorded from sterile electrode needles (29-gauge) inserted subcutaneously into the right upper chest with the negative electrode needle and into the left bottom chest with the positive electrode needle. The signal was then acquired and analyzed using a digital acquisition and analysis system (Power Lab; AD Instruments; LabChart 8 Pro software version). ECG parameters (QT interval) were quantified after 1–2 min from the stabilized trace.

### Histology

Mouse hearts were dissected, rinsed briefly in saline, and fixed overnight in 10% neutral buffered formalin. Following fixation, tissues were dehydrated in 70% ethanol, and submitted to HistoWiz Inc. (Long Island City NY) for sectioning and staining using a standardized automated workflow. Serial transverse sections (5µm) were stained with hematoxylin and eosin (H&E) or Masson’s trichrome. Whole-slide scanning (40×) was performed on an Aperio AT2 (Leica Biosystems).

### *Ex vivo* tissue processing

Mice were anesthetized with 3-4% isoflurane and euthanized via cervical dislocation, followed by heart and/or liver tissue collection. Whole hearts were harvested and cut to small pieces. Per liver, one piece was taken from the left median lobe. Tissue was placed in 1.5 mL tubes containing metal beads (Next Advance Inc., NAVYR1RNA) and flash-frozen in liquid nitrogen. Tissue homogenization was performed using Precellys Evolution Touch Homogenizer (Bertin Technologies P002511-PEVT0-A.0) with 5 cycles of 20 s 6500 rpm shaking and 10 s pause.

For DNA and RNA extraction from the same heart sample, tissue was homogenized in QIAzol Lysis Reagent (Qiagen 79306) and processed using Direct-zol DNA/RNA Miniprep kit (Zymo R2080). For DNA extraction only, heart tissue was homogenized in extraction buffer (Extraction Buffer (Abcam ab193970) + Extraction Enhancer (Abcam ab193971) + cOmplete Protease Inhibitor Cocktail (Roche 11697498001)) and digested with proteinase K (Thermo Scientific EO0491) at 55°C for 16 h, followed by 95°C for 20 min, and gDNA was amplified directly from lysate.

Heart RNA was treated with TURBO DNase (Thermo Fisher Scientific AM2238) at 37°C for 45 min then purified with *Quick*-RNA Miniprep kit (Zymo R1054). Treated RNA was reverse transcribed using SuperScript IV First-Strand Synthesis System (Thermo Fisher Scientific 18091050) with random hexamers.

Liver tissue was processed using Quick-DNA Miniprep Plus Kit (Zymo D4068). Protocol was altered slightly to include a homogenization step after addition of 400 μl Zymo kit lysis buffer (solid tissues buffer and water) to tissue. After homogenization, samples were digested with proteinase K for 1 hr at 55°C (1000 rpm shaking) and then centrifuged at 6000 g for 30 s. Extraction was continued according to manufacturer’s protocol using 200 μl supernatant.

### Quantitative real-time PCR analysis

qPCR reactions for splicing-related genes (*Ttn* and *Camk2d*) and for transgene (PE-N and PE-C cassettes) were performed using PowerUp SYBR Green Master Mix (Applied Biosystems) on a QuantStudio 7 Real-Time PCR System (Applied Biosystems). Primer sequences were as follows: *Ttn N2B* forward: [GGAGTACACCTGCAAAGCCT], reverse: [TGCGGCTTAGGTTCAGGAAG]; *Ttn N2BA* forward: [GGAGTACACCTGCAAAGCCT], reverse: [CCTTGGGCCTGGAGAGAAAG] *Camk2d-A* forward: [GCTCTACTGTTGCCTCCATG], reverse: [GCTGGTTACCACGTTGGC]; *Camk2d-B* forward: [GTCCAGTTCGAGTGTTCAGA], reverse: [GCAGTGAGGCCTGGATCAC]; *Camk2d-C* forward: [GCAGATGGGGTAAAGGAGT], reverse: [GGCAGTGAGGCCTGGATCAC]; PE-N forward: [CGGATCTGCTATCTGCAAGAGATC], reverse: [CTCGTGCTTCTTATCCTCTTCCAC]; PE-C forward: [TGGTGGTGGCCAAAGTGGAAAAG], reverse: [AGTCGATGGGATTCTTCTCGAAGC]. For heart failure and cardiac fibrosis-related genes, qPCR was performed using TaqMan Fast Advanced Master Mix (Applied Biosystems) with the following TaqMan Gene Expression Assays: *Gapdh* (Mm99999915_g1); *Nppa* (Mm01255747_g1); *Nppb* (Mm01255770_g1); *Tgfb1* (Mm01178820_m1); *Tgfb2* (Mm00436955_m1); *Col1a1* (Mm00801666_g1); *Col3a1* (Mm00802300_m1)

Gene expression levels were normalized to *Gapdh* and reported as fold-change relative to the control group.

### Next-generation sequencing and data analysis

Genomic loci of interest were amplified from gRNA or cDNA via two rounds of PCR with CloneAmp HiFi PCR Premix (Takara 639298). The initial PCR step was done using primers to amplify the genomic locus and incorporate the primer binding site for the second round of PCR. PCR reactions were amplified with the following conditions: 98°C for 30 s; n cycles of 98°C for 10 s, 60°C for 5 s and 72°C for 30 s (n = 16 when amplifying cDNA; n = 21-25 when amplifying gDNA). The following primer pairs were used in the initial PCR step to amplify the specified loci:

Mouse *Rbm20* editing site gDNA:

[tcgtcggcagcgtcagatgtgtataagagacagTCTCTTCCCTTCCTCCCAGGTATG] and [gtctcgtgggctcggagatgtgtataagagacagTGTGCCCAAACATCCAACCTGTC]

Humanized mouse *RBM20* DNA editing site gDNA:

[tcgtcggcagcgtcagatgtgtataagagacagCTCCTTGGCTCCCTCACAGATATG] and [gtctcgtgggctcggagatgtgtataagagacagTGTGCCCAAACATCCAACCTGTC]

Mouse *Rbm20* and humanized mouse *RBM20* editing site cDNA:

[tcgtcggcagcgtcagatgtgtataagagacagGACATGCTCCGGGAAGCTGAC] and [gtctcgtgggctcggagatgtgtataagagacagTGTGCCCAAACATCCAACCTGTC]

Human *RBM20* editing site gDNA:

[tcgtcggcagcgtcagatgtgtataagagacagCTCCTTGGCTCCCTCACAGATATG] and [gtctcgtgggctcggagatgtgtataagagacagTCCTGGCATAGGGAGAGTGCTC]

Human *RNF2* editing site gDNA:

[tcgtcggcagcgtcagatgtgtataagagacagAGGCTGTGCAGACAAACGGAAC] and [gtctcgtgggctcggagatgtgtataagagacagCCAACATACAGAAGTCAGGAATGC]

Mouse *Dnmt1* editing site gDNA:

[tcgtcggcagcgtcagatgtgtataagagacagGGAGGCAAGCGCAGGCACTC] and [gtctcgtgggctcggagatgtgtataagagacagTTGCCCTGTGTGGTACATGCTGC]

Mouse PE self-inactivation site:

[tcgtcggcagcgtcagatgtgtataagagacagAGACGCCTCCAGGCCACCATG] and [gtctcgtgggctcggagatgtgtataagagacagTTCTTGCTGGGCACCTTGTACTC]

Unique Illumina index 1 and index 2 adapters were incorporated during the second PCR step by amplifying first-round PCR product with Illumina DNA/RNA UD Indexes (Illumina 20091646, 20091647, 20061648, and 20061649). PCR reactions were amplified with the following cycle conditions: 98°C for 30 s; 9 cycles of 98°C for 10 s, 55°C for 5 s, and 72°C for 30 s. Second-round PCR product was size-selected and purified using E-Gel SizeSelect II Agarose Gels, 2% (Invitrogen G661012). Purified libraries were quantified using Qubit dsDNA HS assay kit (Thermo Fisher Q33231) and pooled for sequencing on Illumina NextSeq 550 using either the NextSeq 500/550 High Output Reagent v2.5 75 cycles kit (20024906) or 150 cycles kit (20024907), or, for sequencing of *in vitro* libraries, using the NextSeq 500/550 Mid Output v2.5 150 cycles kit (20024904).

Sequencing reads were de-multiplexed through Illumina’s BaseSpace Sequence Hub. Sequence extraction, unique sequence counting, and editing analysis were performed using an in-house Python workflow. Briefly, a sequence of 36bp covering the editing site was extracted from NGS reads with Q score > 30 (or > 20 for *in vitro* samples) at all positions in the extracted region. Then, unique sequences were counted and sequence counts of each test sample were compared against that of a control to generate a list of sequences occurring in the test sample either above a frequency of 0.01 or enriched 3-fold above that sequence in the control and occurring in the test sample above a frequency of 3 in a million (1 in ten thousands for *in vitro* samples). This step removes most sequence variants generated by PCR errors and NGS errors. Finally, the frequency of various PE outcomes (i.e. WT sequence, mutant sequence, fully edited sequence, partially edited sequences, mutations and indels) were calculated from that list.

### Statistical analysis

All data are shown as mean ± SD, mean ± SEM, mean ± SD with individual values from biological replicates, or mean ± SEM with individual values from biological replicates as indicated in the figure legends. Statistical analyses were performed with GraphPad Prism software. Statistical methods are indicated in the figure legends.

## List of Supplementary Materials

Fig. S1 to S6

## Acknowledgements

We thank Eric Olson and his laboratory for providing the *Rbm20^R636Q^* mouse model. We thank the vivarium team at Tenaya Therapeutics for animal husbandry support.

## Author contributions

W.L. generated the humanized *RBM20* mouse model and conducted *in vivo* experiments and *ex vivo* analyses. L.M.R. cloned PE constructs, manufactured AAV for *in vivo* experiments, and conducted *ex vivo* analyses. E.E. developed guide RNAs targeting human *RBM20*, cloned PE constructs, and manufactured AAV for *in vivo* experiments. H.Z. designed the humanized *RBM20* mouse model and performed MEF isolation. C.D.H. contributed to mouse genotyping and *in vivo* experiments. A.G.S., T.H., L.M.L., and K.N.I. supervised the research. Z.C. designed the studies and performed viral injections. W.L., L.M.R., and Z.C. prepared the manuscript with input from all authors.

## Data and materials availability

All data supporting the findings described in this manuscript are available in the article and in the Supplementary Materials. Sequences are available in the Materials and Methods and Supplementary Materials.

## Supplementary Materials

**Fig. S1.**
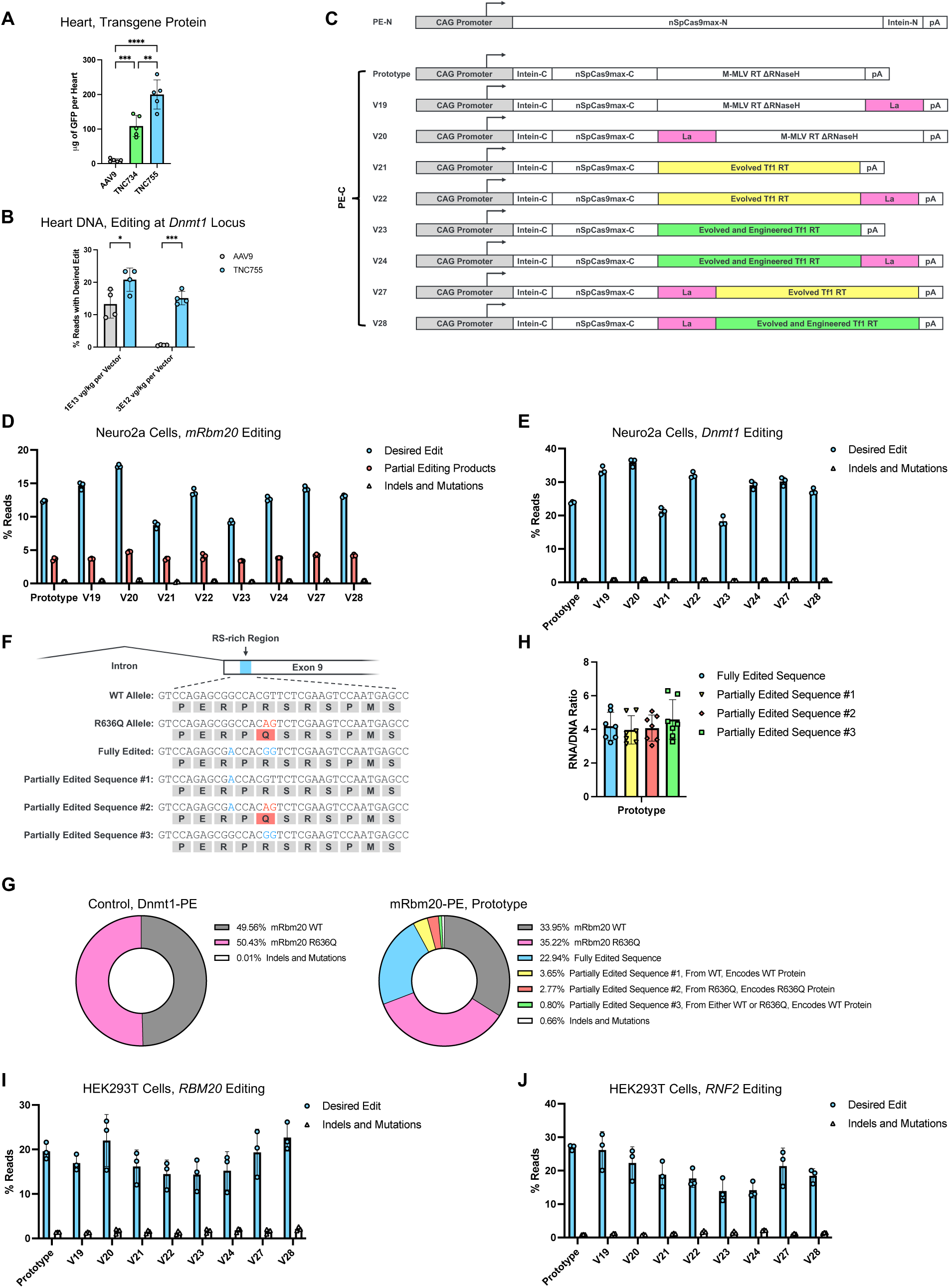
Developing a dual-AAV based strategy for *in vivo* cardiac prime editing (PE). (**A**) Expression levels of EGFP transgene protein in the heart mediated by AAV9 and in-house engineered capsids. C57BL/6J mice at 6 weeks of age were administered with either WT AAV9 or one of our in-house engineered capsids (sequences available upon request) containing EGFP transgene under control of the 400bp *TNNT2*-derived promoter (1E13 vg/kg, 5 animals per group) via RO injection. Mice were euthanized 3 weeks post-injection and EGFP expression in the heart was assessed using Abcam GFP ELISA kit (ab171581). (**B**) *In vivo* cardiac editing efficiencies and cardiac transduction levels mediated by AAV9 and TNC755. TNC755 was used in efficacy studies described in this manuscript. C57BL/6J mice were administered with either WT AAV9 or one of our engineered capsids containing PE-N and PE-C cassettes targeting the *Dnmt1* locus (1E13 vg/kg or 3E12 vg/kg, 4 animals per group) via RO injection. Mice were euthanized 3 weeks post-injection and DNA and RNA were isolated from whole heart. *Dnmt1* editing efficiency was determined via NGS and transgene DNA transduction and RNA expression were quantified via qPCR (normalized to GAPDH, average of PE-N and PE-C). For A and B, dots represent individual mice. Statistical significance was calculated using one-way ANOVA with Tukey’s multiple comparisons test. \**P* < 0.05, \*\**P* < 0.01, \*\*\**P* < 0.001, \*\*\*\**P* < 0.0001. (**C**) Schematic of our PE cassettes for *in vitro* experiments. (**D** and **E**) *In vitro* editing efficiency at the target locus of Neuro2a cells transfected with PE-N and PE-C cassettes and pegRNA/ngRNA targeting either the mouse *Rbm20* (D) or *Dnmt1* (E) locus. (**F**) Schematic demonstrating possible post-edit outcomes in *Rbm20^R636Q/+^* mice edited using our mRbm20-PE. (**G**) Mean percentages of *Rbm20* editing outcomes in *Rbm20^R636Q/+^*mice treated with Dnmt1-PE (control) or mRbm20-PE prototype. (**H**) RNA/DNA ratio of different post-edit sequences from mice treated with mRbm20-PE Prototype. Dots represent individual mice and error bars represent mean ± SD. (**I** and **J**) *In vitro* editing efficiency at the target locus of HEK293T cells transfected with PE-N and PE-C cassettes and pegRNA/ngRNA targeting either the *RBM20* (I) or *RNF2* (J) locus. For D, E, I, and J, dots represent individual biological replicates, and error bars represent mean ± SD.

**Fig. S2.**
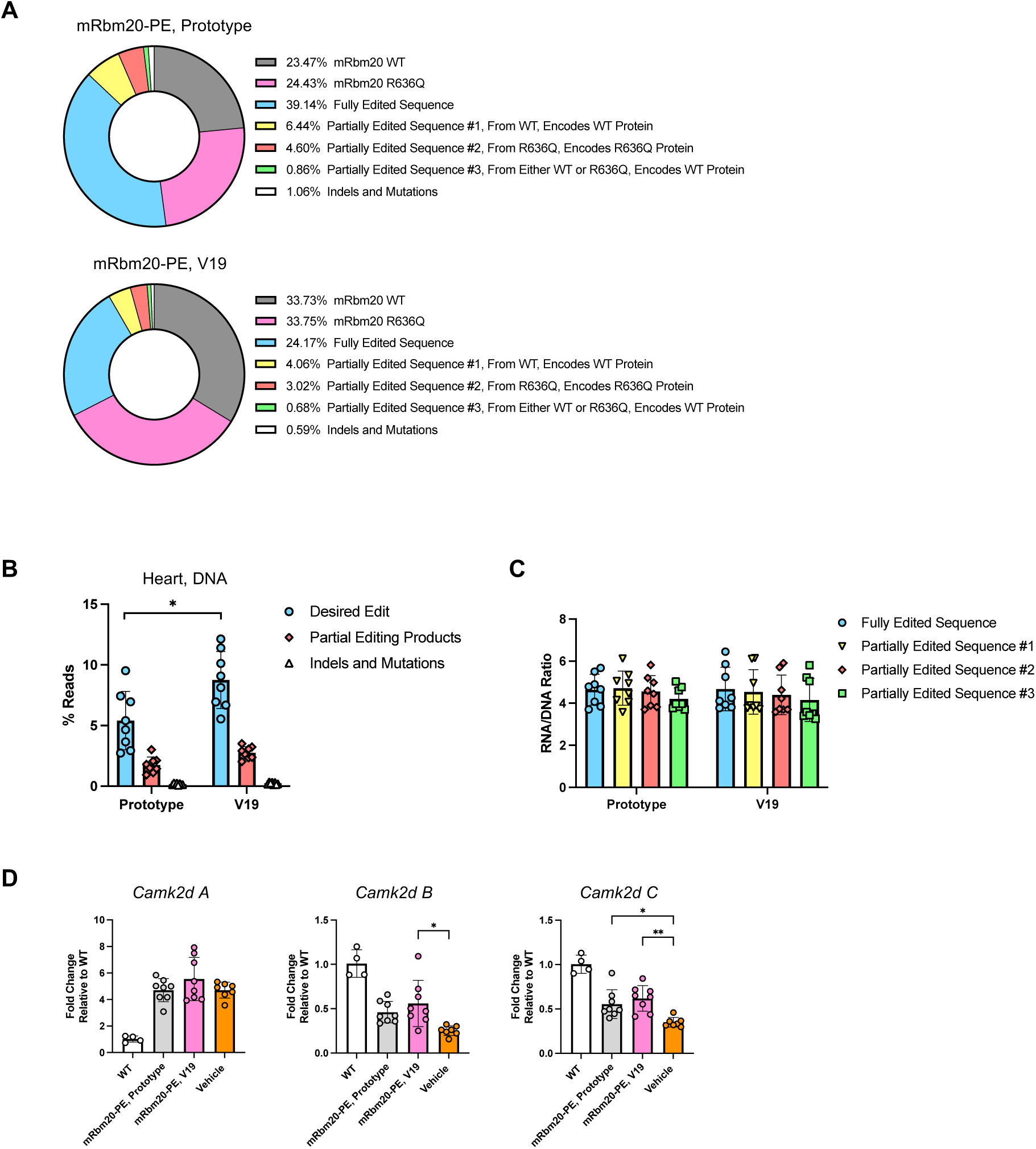
V19 prime editor improves editing efficiency and rescues phenotypes in *Rbm20^R636Q^* DCM mouse model. (**A**) Mean percentages of *Rbm20* editing outcomes in *Rbm20^R636Q/+^* mice treated with mRbm20-PE Prototype or mRbm20-PE V19 (4.6E13 vg/kg per vector). (**B**) Editing efficiency at the *Rbm20* locus, detected from whole heart DNA samples. (**C**) RNA/DNA ratio of different post-edit sequences. (**D**) Splicing patterns of *Camk2d* gene. For B to D, dots represent individual mice, error bars represent mean ± SD. For B, statistical significance was calculated using unpaired t-test. For D, statistical significance was calculated using two-way ANOVA with Tukey’s multiple comparisons test. \**P* < 0.05, \*\**P* < 0.01.

**Fig. S3.**
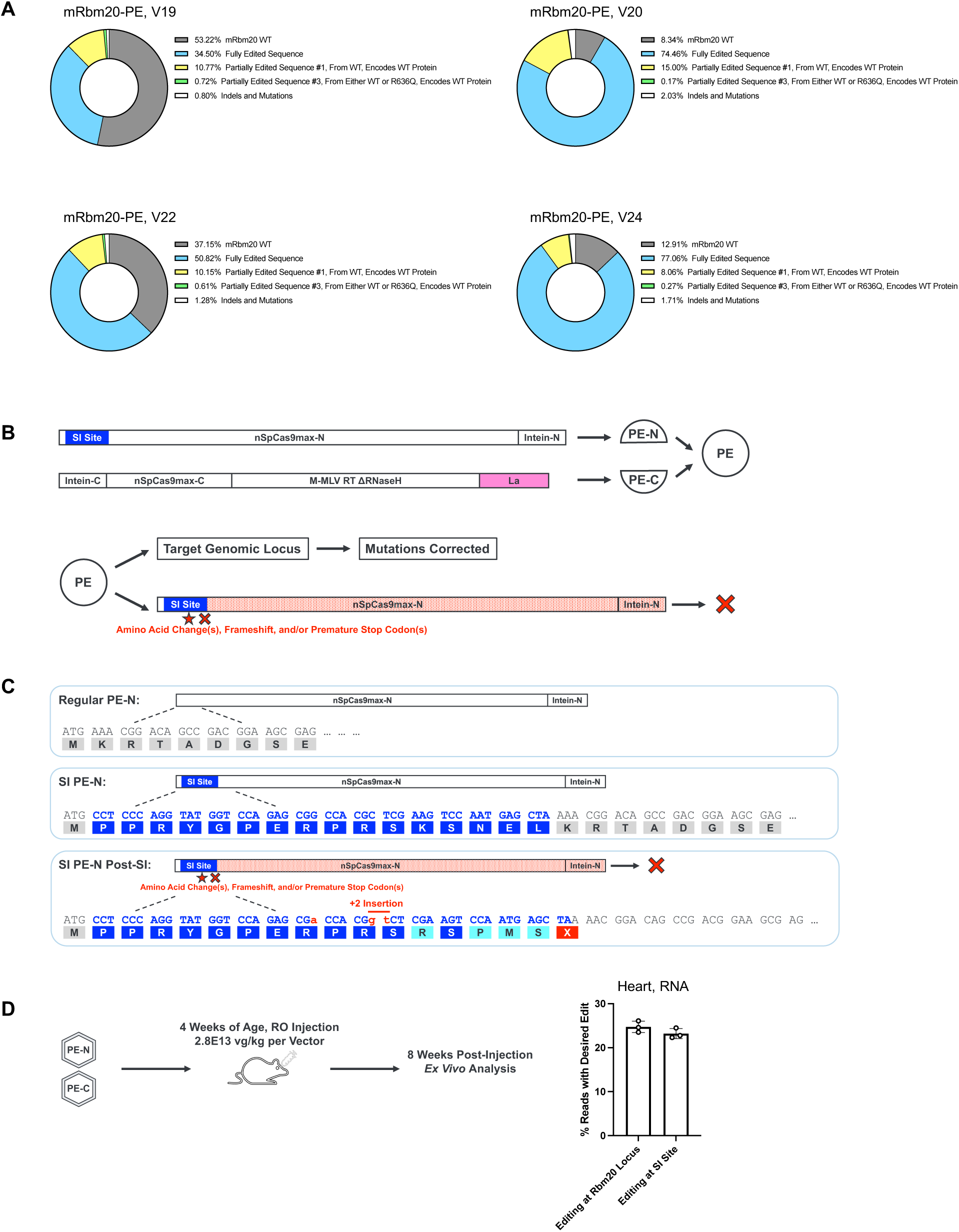
Engineering prime editors to enhance editing efficiency and/or enable self-inactivation. (**A**) Mean percentages of *Rbm20* editing outcomes in mice treated with mRbm20-PE V19, V20, V22, and V24 (2.E13 vg/kg per vector). WT mice were used in this study. (**B**) Schematic detailing our self-inactivation (SI) strategy: an SI site is a polynucleotide sequence which encodes a short peptide and can be recognized and edited by the prime editor; the SI site is incorporated to the coding region of PE-N and/or PE-C; the unedited SI site adds a short peptide to the PE protein and the resulting fusion protein is still functional; once introduced to the cell, PE happens at both the genomic target locus and the SI site; PE at the SI site introduces amino acid change(s), frameshift, and/or premature stop codon(s); the post-editing SI site inactivates the expression and/or function of the PE protein. (**C**) Schematic of a SI PE example, carrying an SI site that can be edited by our mouse *Rbm20*-targeting pegRNA. (**D**) *In vivo* cardiac editing at *Rbm20* locus and SI site by mRBM20-SI-PE V19. *Ex vivo* analysis was performed at 8 weeks post-injection. Dots represent individual mice, and error bars represent mean ± SD.

**Fig. S4.**
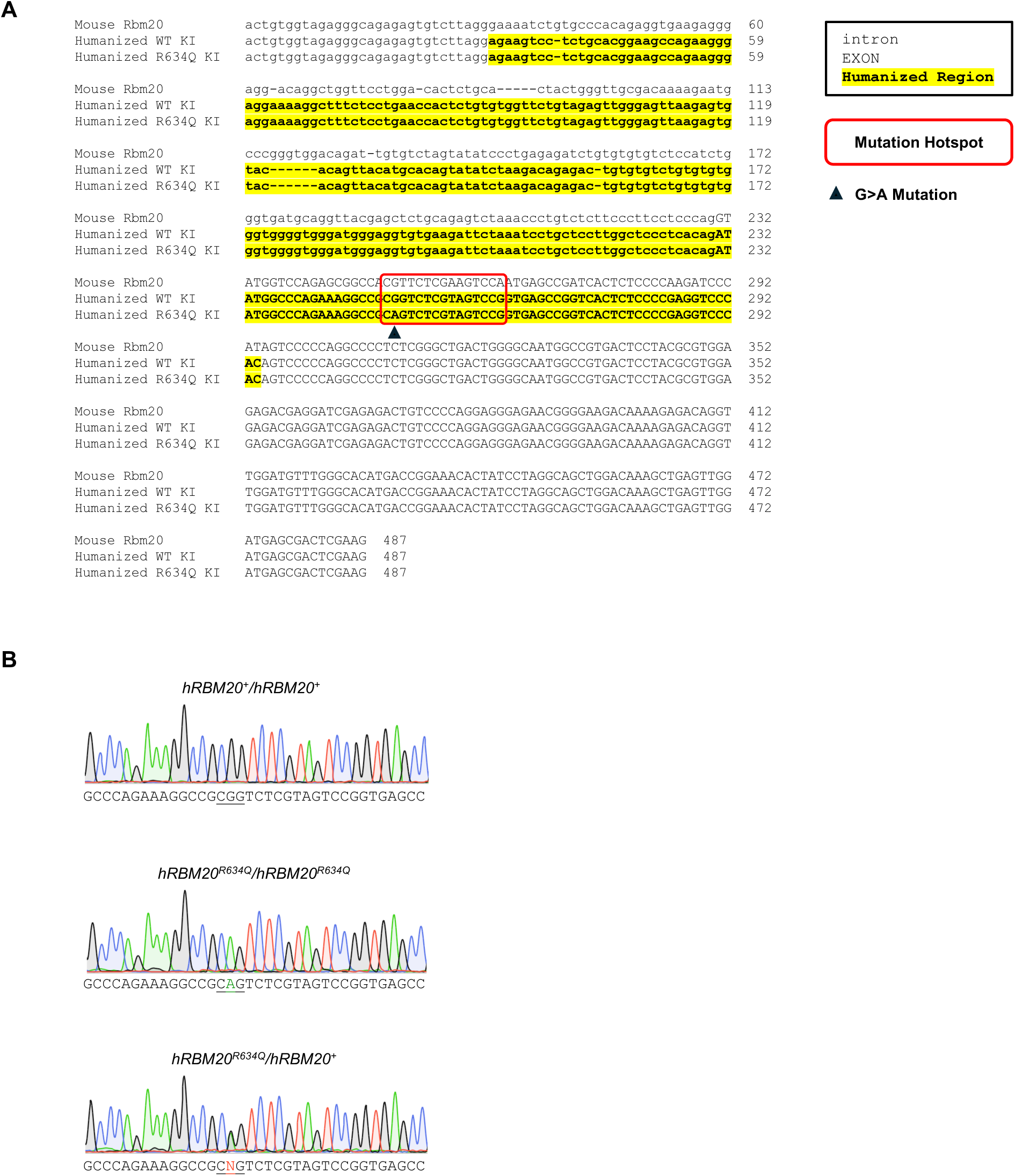
Generating a novel humanized *RBM20* mouse model. (**A**) Sequence comparison of mouse *Rbm20*, human *RBM20*, and human *RBM20^R634Q^* indicating intron and exon regions, humanized region, mutation hotspot, and R634Q G>A mutation. (**B**) Sanger sequencing confirmation of humanized mouse genotypes.

**Fig. S5.**
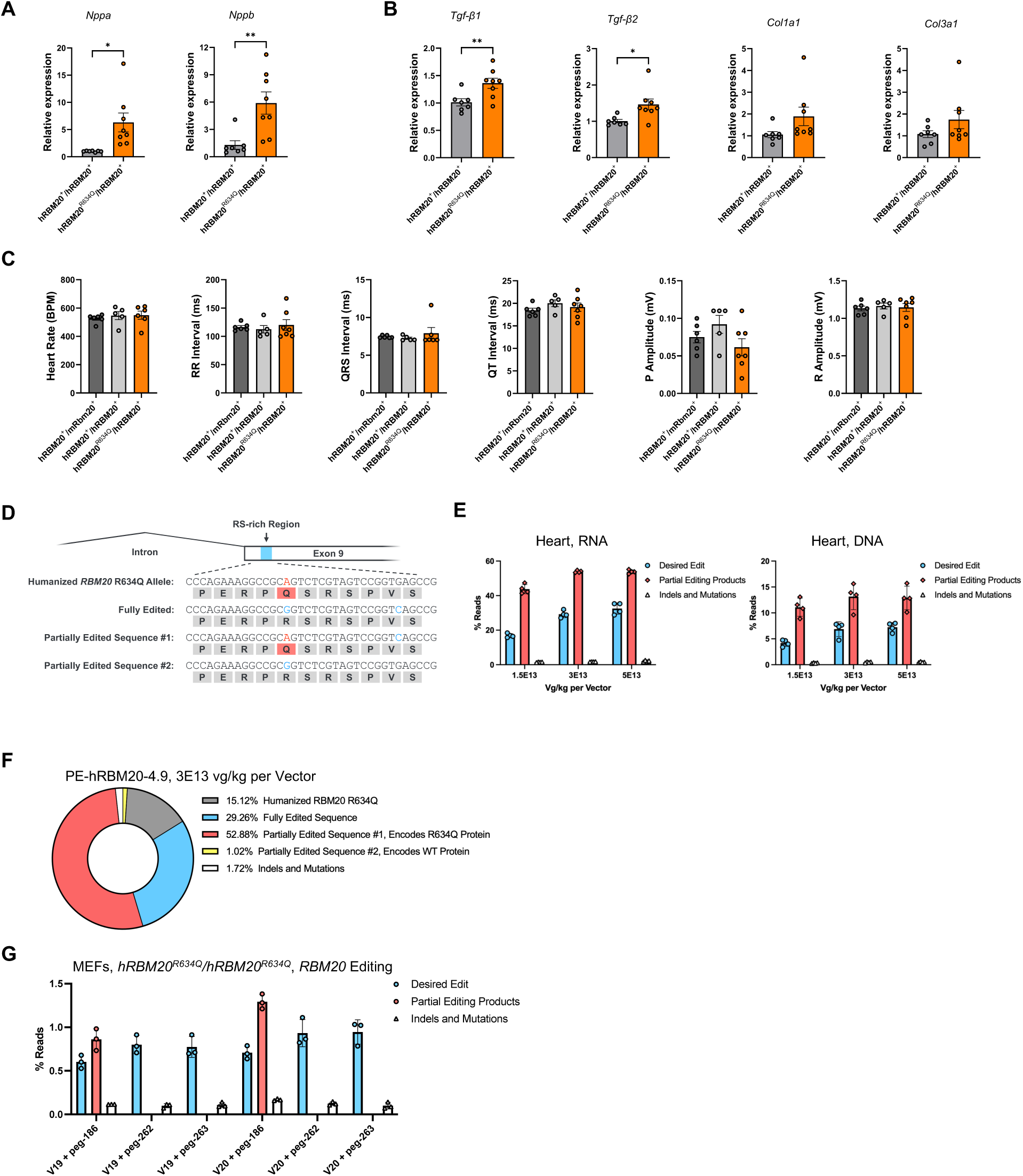
Humanized *RBM20* mouse model exhibits *RBM20* cardiomyopathy phenotypes and facilitates the development of human *RBM20* PE. Quantitative PCR analysis of (**A**) heart failure markers *Nppa* and *Nppb* and (**B**) fibrosis-related genes in heterozygous mouse hearts. (**C**) Electrocardiogram analysis showed normal electrophysiological parameters in heterozygous mice. (**D**) Schematic demonstrating possible post-edit outcomes for *hRBM20^R634Q^* allele edited using PE-hRBM20-4.9. (**E**) *hRBM20^R634Q^* editing outcomes in RNA or DNA isolated from whole heart of *hRBM20^R634Q^/mRBM20^+^*mice treated with PE-hRBM20-4.9 at 1.5E13, 3E13, or 5E13 vg/kg per vector. (**F**) Mean percentage of *RBM20* editing outcomes in RNA isolated from whole heart of *hRBM20^R634Q^/mRBM20^+^* mice treated with PE-hRBM20-4.9 at 3E13 vg/kg per vector. Dots represent individual mice, error bars represent mean ± SD. \**P* < 0.05, \*\**P* < 0.01. (**G**) *In vitro RBM20* editing efficiency of *hRBM20^R634Q^/hRBM20^R634Q^* MEF cells transfected with *in vitro* V19 or V20 PE cassette containing h-ng-26 and one of three pegRNAs. Dots represent biological replicates, error bars represent mean ± SD.

**Fig. S6.**
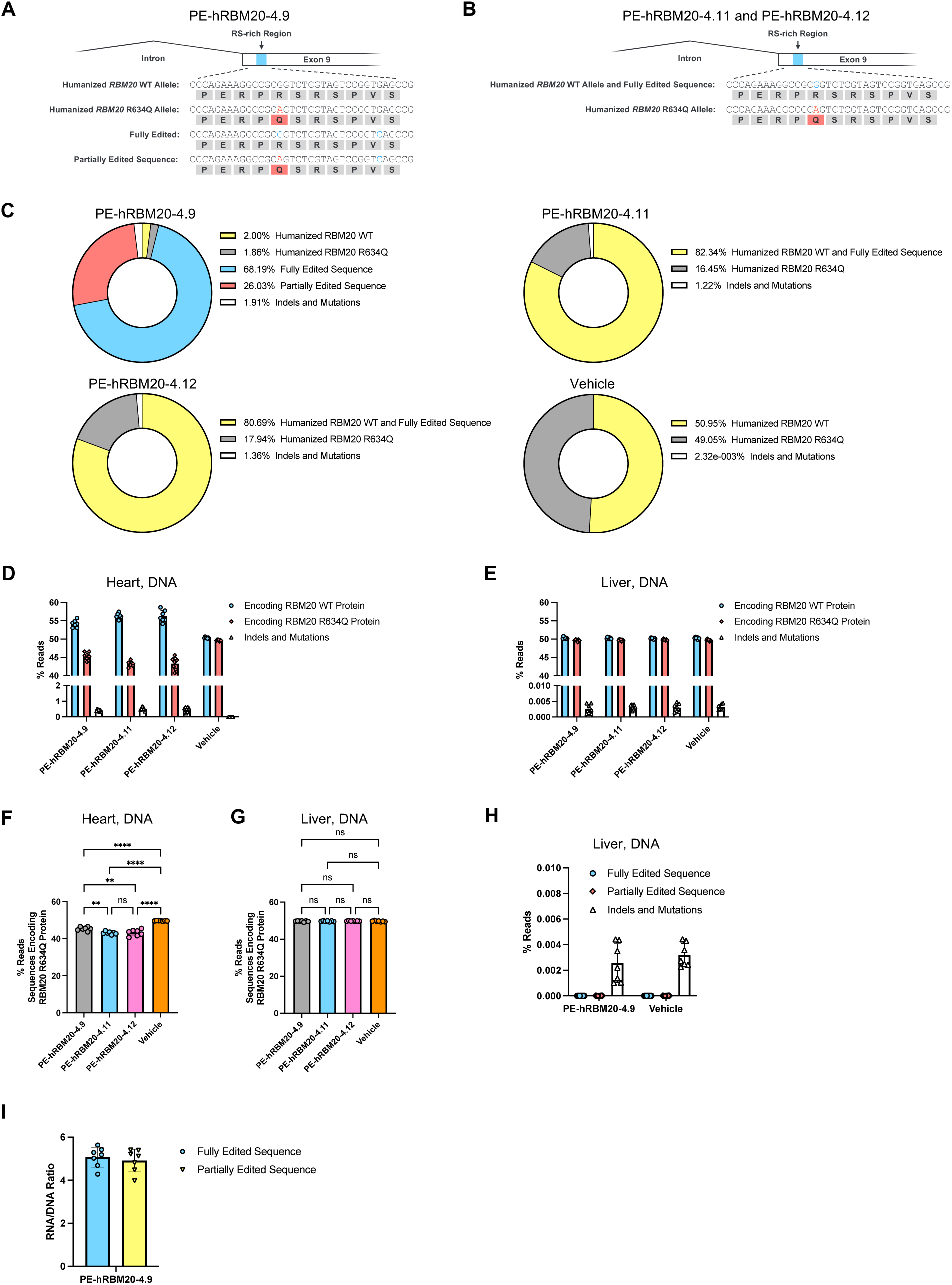
*In vivo* cardiac PE successfully corrects *RBM20^R634Q^* in mouse heart without showing detectable editing in off-target tissue. (**A** and **B**) Schematic depicting editing outcomes of humanized *hRBM20^R634Q^/hRBM20^+^*mice treated with PE-hRBM20-4.9 (A) or PE-hRBM20-4.11 and PE-hRBM20-4.12 (B). (**C**) Mean percentages of *RBM20* editing outcomes in RNA of *hRBM20^R634Q^/hRBM20^+^* mice treated with PE-hRBM20-4.9, PE-hRBM20-4.11, PE-hRBM20-4.12, and vehicle. (**D** and **F**) Editing efficiency at humanized *RBM20* locus, detected from whole heart DNA samples. (**E**, **G**, and **H**) Editing efficiency at humanized *RBM20* locus, detected from left median lobe liver DNA samples. (**I**) RNA/DNA ratio of fully edited sequence and partially edited sequence outcomes for humanized *hRBM20^R634Q^/hRBM20^+^*mice treated with PE-hRBM20-4.9. For D to E, each dot represents one animal. For F and G, statistical significance was calculated using one-way ANOVA with Tukey’s multiple comparisons test. \*\**P* < 0.01, \*\*\*\**P* < 0.0001.

